# TBDB – A database of structurally annotated T-box riboswitch:tRNA pairs

**DOI:** 10.1101/2020.06.17.157016

**Authors:** Jorge A. Marchand, Merrick D. Pierson Smela, Thomas H. H. Jordan, Kamesh Narasimhan, George M. Church

## Abstract

T-box riboswitches constitute a large family of tRNA-binding leader sequences that play a central role in gene regulation in many gram-positive bacteria. Accurate inference of the tRNA binding to T-boxes is critical to predict their cis-regulatory activity. However, there is no central repository of information on the tRNA binding specificities of T-box riboswitches and *de novo* prediction of binding specificities requires advance knowledge of computational tools to annotate riboswitch secondary structure features. Here we present **T-box** annotation **D**ata**b**ase (**TBDB,** https://tbdb.io), an open-access database with a collection of 23,497 T-box sequences, spanning the major phyla of 3,621 bacterial species. Among structural predictions, the TBDB also identifies specifier sequences, cognate tRNA binding partners, and downstream regulatory target. To our knowledge, the TBDB presents the largest collection of feature, sequence, and structural annotations carried out on this important family of regulatory RNA.

## Introduction

Bacteria exploit a wide-range of *cis*-acting RNA regulatory elements to control gene expression in response to specific environmental stimuli. One strategy used for modulating gene expression involves using 5′-UTR leader riboswitches to regulate transcription or translation [1–3]. The transcriptional or translational logic of riboswitch leader sequences are conditionally dependent on the binding of a specific ligand [4].

In the gram-positive model organism *Bacillus subtilis,* an analysis of *cis*-regulatory sequences in the upstream region of several amino-acyl tRNA synthetase (ARS) genes revealed that non-amino-acylated tRNAs can act as a positive regulator [5,6]. The discovery of this feedback loop was a breakthrough in understanding the expression of ARSs under nutrient limiting conditions. The T-box leader sequence was the first classical “riboswitch” family to be discovered, preceding the discovering of metabolite-binding ribo-regulators [6].

T-box leader sequences are *cis*-acting RNA elements that link either transcription or translation of downstream genes to the aminoacylation-state of tRNA [7–9]. In transcriptional regulation, the 3′-end of T-box leader sequences folds into either a terminator structure, prematurely stopping transcription, or antiterminator structure, allowing transcription to proceed (**Figure 1A**). Translational regulation occurs through a similar two-state mechanism, whereby a ribosome-binding site is either structurally sequestered, preventing ribosome binding, or exposed, allowing ribosome binding and translation [8].

**Fig 1.**
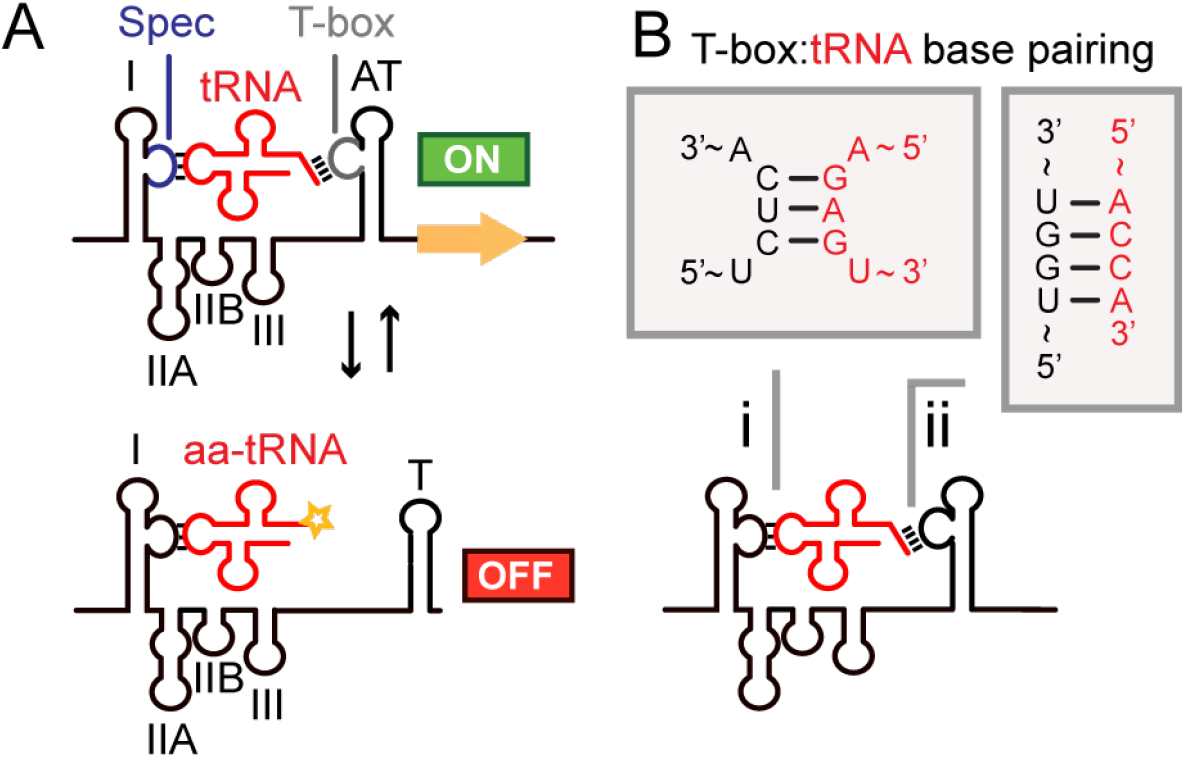
T-box riboswitches are *cis*-regulatory elements that use tRNA as a ligand. T-boxes control translation or transcription of downstream genes. When uncharged cognate tRNA binds the T-box, transcription or translation can proceed through stabilization of the antiterminator/antisequestrator structures. If charged cognate tRNA binds the T-box, a terminator/sequestrator secondary structure forms preventing transcription or translation of downstream gene. A) An archetypal “two-state” conformational switch of a transcriptional T-box is shown with structural features labeled: I = stem I; IIA = stem IIA; IIB = stem IIB; III = stem III; AT = antiterminator; T = terminator; Spec = specifier loop (blue); T-box = T-box sequence (gray). B) Watson-Crick base pairing between T-box and tRNA in two critical regions dictate T-box specificity: (i) Specifier:anticodon base pairing dictates tRNA ligand binding while (ii) T-box:acceptor arm base pairing controls regulatory logic.

Mechanistic studies on T-box riboswitches have revealed important interactions between T-boxes and tRNAs. Classical Watson-Crick base pairing has been shown to occur between the T-box specifier sequence and the tRNA anticodon [10]. Additional Watson-Crick base pair between the tRNA acceptor arm (‘5-NCCA-3′) and the ‘T-box sequence’ (5′-UGGN-3′) was demonstrated to control regulatory logic (**Figure 1B**) [11,12]. These interactions confer initial T-box specificity to its cognate tRNA. If un-charged tRNA binds to the T-box, Watson-Crick base pairing between the ‘T-box sequence’ and acceptor arm causes the T-box leader to adopt an antiterminator fold (for transcriptional regulation) or antisequestrator fold (for translational regulation). If the bound species is charged-tRNA either a steric barrier or a charged tRNA dependent conformational change prevents acceptor arm base pairing, leading the T-box to adopt a terminator fold (for transcriptional regulation) or sequestrator fold (for translational regulation).

Structural studies of T-box:tRNA complexes have highlighted the importance of additional interactions between T-boxes and cognate tRNAs. Initial X-ray crystallographic studies of Stem I complexed with tRNA provided evidence of π-stacking interactions between the apical portion of Stem I and tRNA D-loop [2,11,13]. More recent X-ray and cryoEM structures of the full complex show several structural folds of the T-box including the variable regions namely, Stem IIA, and Stem IIB engaging the tRNA in specific binding interactions [14–17]. These structural findings are supported by *in vitro* transcription studies where it was shown that Stem II contacts with cognate tRNA are important for tuning binding between a *Clostridium* sp. alanyl T-box and alanyl-tRNA [18].

The intricate and specific interactions between the T-box:tRNA pair can be leveraged for a variety of applications across basic research and bioengineering. For example, a recent study used a glyQS T-box to engineer a ribozyme that can specifically charge tRNA^gly^ for use in cell-free protein synthesis [19]. T-boxes also have the potential to be used as a generalizable ‘registry-of-parts’, capable of independently sensing amino acid levels in the environment with a single RNA element[20]. Furthermore, due to their prevalence and importance in gram-positive bacteria, T-boxes are also being studied as targets for antibiotic therapeutics [21,22]. Bacterial genomes tend to have several hypothetical proteins, uncharacterized genes with remote homologs or amino-acid transporters whose functions cannot be reliably predicted from sequence similarity alone. In this regard T-box specifier prediction has been used as a tool to uncover the function of unknown *cis*-regulated genes [23,24]. In one case, the predicted T-box family was used to infer the substrate specificity of downstream amino acid transporters [25].

Despite the identification of several thousand leader sequences across various databases, T-box structures and functions remain under characterized [23–30]. Existing public databases which host putative T-box sequences do not include potential tRNA partners for annotated T-boxes nor predict the tRNA binding specificities. Currently, *in silico* structure prediction and feature extraction is required to predict tRNA substrates from raw sequences, and therefore exists as a barrier for entry to anyone interested in T-box research.

Here we present the **T-box** annotation **D**ata**b**ase (**TBDB**), a compilation of T-box sequences from various primary sources with detailed annotations to aid future research. TBDB predicts putative transcriptional and translational T-box sequences, annotates secondary structures, identifies functional features and downstream genes, finds cognate pairs of tRNAs from host organisms, calculates MFE (minimum free-energy) structures, and provides rich visualization for known and predicted T-box leader sequences (**Figure 2**). The TBDB is browsable at https://tbdb.io, with the whole database available to download as a single flat file. The TBDB will be a valuable resource for studying canonical, engineered, and mutant T-box mechanisms and will provide a point-of-entry for studying regulation and interactions between T-box:tRNA pairs. As a resource for the wider non-coding RNA community, the TBDB is the first structural and functionally annotated database for studying gene regulation by T-box riboswitches.

**Fig 2.**
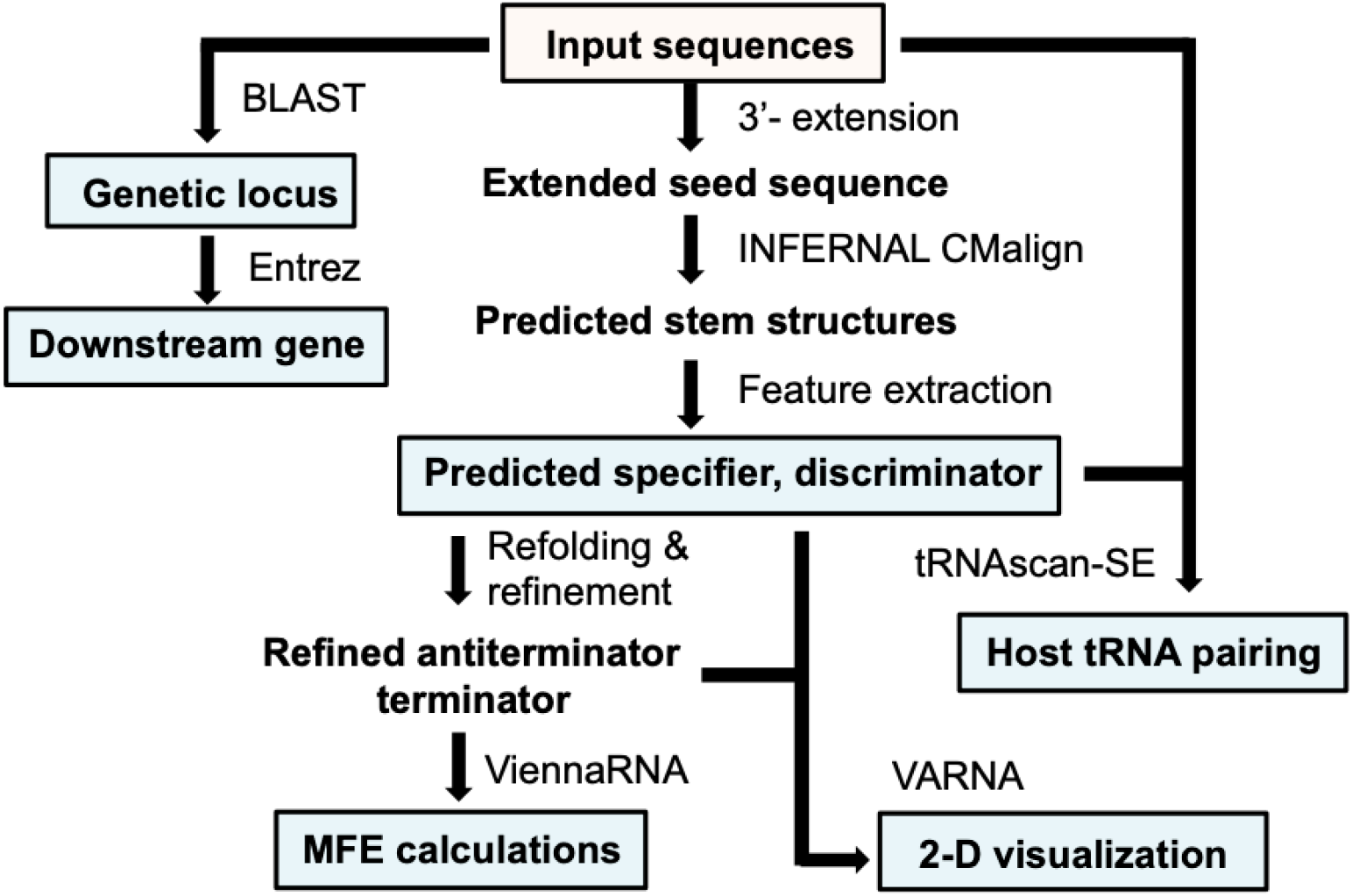
Flow diagram - Construction of TBDB. T-box structures were predicted from input sequences using INFERNAL and RNAfold (Vienna RNA) [33,34]. T-box features (specifier sequence and T-box sequence) were extracted from structural predictions. For input sequences where genomic information was not provided, BLAST (NCBI) was used to identify genetic locus within host. Entrez (NCBI) queries were used to compile all genomic sequence records of T-box host organisms including genes found downstream of T-box input sequences. tRNAscan-SE was run on all genomes to find tRNAs in the hosts with antiterminator sequences that are reverse complements of T-box specifier sequences. Predicted, structures were refined using ViennaRNA. Refined structures, with predicted features, were visualized as 2-D representations using VARNA. Minimum free energy (MFE) calculations were performed using ViennaRNA on refined terminator and antiterminator structures.

## Results and Discussion

### Accessing TBDB content

The TBDB aims to be a comprehensive and approachable hub for predictions of T-box structure and function. Database entries are provided in a searchable, tabulated format. Users can query entries based on fields that include sequence, host organism, specifier sequence, T-box sequence, or predicted tRNA family **(Figure S1**).

Detailed information on each TBDB entry can be obtained by accessing the unique ID in the database table. Doing so brings users to a T-box entry page that contains source, downstream protein annotation, structural, functional, and sequence information. The title of the entry page provides the unique T-box ID, a unique identifier generated by TBDB, as well as the predicted tRNA family the T-box interacts with, and host organism (**Figure S2**). The source information panel gives a high-level summary of the T-box entry and includes information regarding genetic locus and feature predictions. The following panel provides an interactive genome browser (NCBI) starting at the T-box locus and ending 5000 bp downstream. The genome browser allows users to visualize the genomic context of T-box sequences and provides a quick method to assess the validity of T-box codon specifier predictions. For example, a Trp T-box is observed as a 5′-UTR of an operon involved in tryptophan biosynthesis (**Figure S3**).

Towards the goal of making a T-box registry-of-parts, in the subsequent panel we provide a T-box sequence that stretches from the Stem I to the terminator poly-U region (**Figure S4**). Visualizations (VARNA) of the predicted secondary structures of the T-box are given in the next panel [31]. These 2-D representations highlight important features of the T-box entry including Stem I (light-yellow), specifier bases (dark yellow), antiterminator (light blue), T-box sequence (dark blue), and terminator stem (red). The dot-bracket representations of the 2-D structures also provided (**Figure S5**). Results for tRNA matching, generated using tRNAscan-SE, can be found in the following panel (**Figure S6**) [32]. Here, we show the sequence and description for the highest scoring predicted tRNA, with matching anticodon, in T-box host organism (if available). tRNAs for alternative specifier reading frames are also generated if our model could not identify a consensus specifier. Minimum free energy (MFE) predictions for each of the folds are given in the thermodynamics section, and are the result of structure refinement performed using ViennaRNA (**Figure S7**) [33]. Finally, the INFERNAL panel has output information from structural searches, which can be used as a reference by users interested in structure prediction information and quality (**Figure S8**) [34].

### Sourcing and curation of the TBDB

The T-box annotation Database comprises 23,497 non-redundant T-box sequences. Sequences used to build the TBDB were sourced from databases that either used covariance models or motif analysis for T-box sequence discovery [24,26,35–38]. Duplicate T-boxes were removed as many of the input sequences used to generate the TBDB were shared across the various databases with varying definitions of T-box start and end. We used an INFERNAL covariance model (RFAM RF00230) to predict transcriptional T-box structure from T-box input sequences [34,38]. For translational T-boxes, a custom covariance model was generated using seed sequences for translational T-boxes found in previously published work [8,24].

In order to further curate the database, we only made tRNA predictions for sequences with predicted structures that properly fold into canonical Stem I and antiterminator regions. Proper *in silico* folding of these two regions are critical for identification of specifier sequence and T-box sequence. Sequences for which we could not predict a plausible terminator stem are kept in the database, as they could be members of translational control T-boxes, which do not contain terminators but rather sequestrator stems. In the interest of preserving sequence information, T-boxes with no structural predictions are displayed in TBDB though only genetic context is available as additional information. For transcriptional T-boxes, *in silico* folding of T-box terminator and antiterminator regions were used as an additional metric to examine plausibility of structure predictions (**Figure S9**). The free-energy values for the folds predicted in the antiterminator and terminator regions (Δ*G_anti_*= −7.4 ± 4.2 kcal/mol, Δ*G_term_* = −19.5 ± 7.9 kcal/mol) is largely consistent with the expectation that terminator hairpin is more stable than the antiterminator. In all, the TBDB contains 22,524 putative transcriptional T-boxes and 973 putative translational T-boxes.

### Identification of T-box:tRNA pairs

T-boxes tend to have a strict preference for canonical Watson-Crick (W-C) base pairing with the anticodons of their cognate tRNA [9]. Deviations from canonical W-C binding in any of the triplet codon positions (the central base being least tolerant, relative to the 5′ and 3′ positions) typically results in impaired tRNA binding and anti-termination [39]. Depending on the length of specifier bulge, it is also likely that there are alternative specifier reading frames, allowing for the possibility of multi-tRNA specificity in gene regulation [40]. However, experimental work uncovering the determinants of multi-specificity in T-boxes remains sparse. Our specifier prediction assignment takes into consideration downstream gene assignment and possible W-C base pairing between the T-box sequence and the tRNA discriminator base from the host organism. In assigning T-box specifier, priority was given to as the region 2-4 bp (inclusive) 5′-from the Stem I specifier bulge, though “−1” and “+1” specifier reading frames were also considered (**Figure S10**). In a majority of the cases, the variable position (5′-UGGN-3′) on the T-box shows W-C base pairing with the discriminator base (5′-**N**CCA-3′) of the cognate tRNA species. Exceptions were noted in the Trp family, where 46% of T-boxes in our collection have a ‘U’ at the degenerate position while their cognate tRNAs have a ‘G’ at the discriminator position, suggesting a G:U wobble pair, as has been previously noted [41]. Based on this observation we allowed for wobble base-pair in our final T-box specifier model for the tRNA discriminator-T-box variable position. In the case of putative His T-boxes, discriminator base matching was not considered as most tRNA^His^ transcripts have an internally paired discriminator base.

In practice, the TBDB identifies the cognate tRNA pairs from T-box hosts by first predicting the specifier sequence, then searching genome records of respective hosts for tRNAs that have a matching anticodon (W-C base-pairing only allowed) and discriminator base-pairing (both W-C and wobble pairing allowed). Our model gave a single specifier frame prediction for 16,010 T-box sequences, two possible specifiers frames for 2,860 sequences, and three specifier frames for 3,651 sequences. For 48 sequences, we were able to predict a specifier but were unable to find the canonical T-box sequence (5′-UGGN-3′). In cases where more than one specifier is possible, preference is given to the “+0” specifier reading frame based on empirical observations of experimentally studied T-boxes (**Figure S10C)**. In all, T-box leaders containing Trp-, Leu-, and Ile-tRNA matching specifiers were most commonly observed in our collection while Lys-, Glu-, and Gln-matching specifiers were the least common. **Supplementary Table 1A & 1B** show composition of the T-box Annotation Database amino acid family and specifier usage.

**Table 1.**
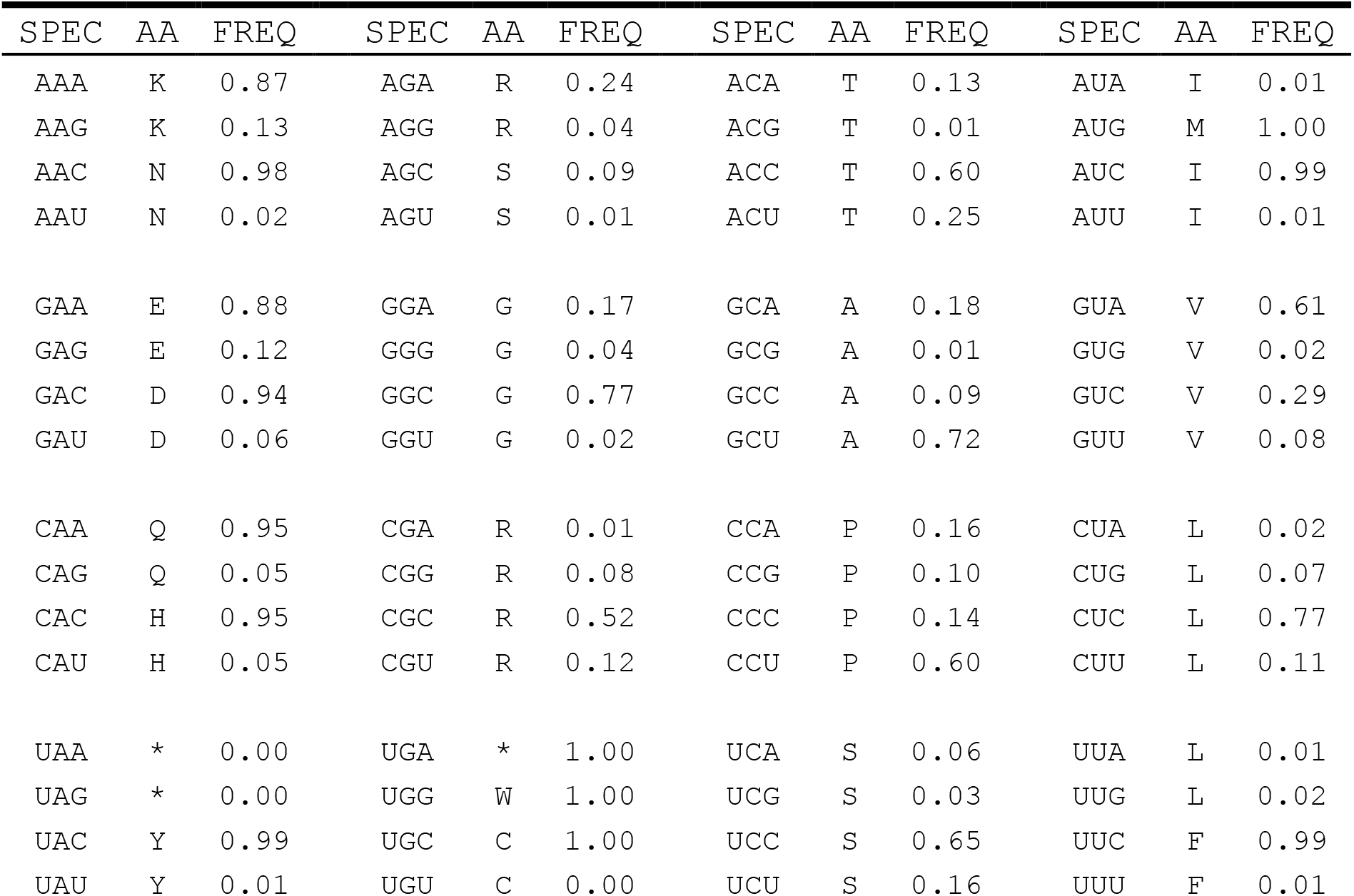
Specifier usage table based on “Top” specifier predictions for T-box leader sequences in the TBDB. Frequency given on an amino acid basis for the entire sequence collection in TBDB.

Through our tRNA search, we were able to match 76.9% of T-box sequences (for which we predicted a specifier) with a tRNA of the native host. Grouping T-boxes by specifier sequence, we found that tRNA-matching was >80% for most specifiers (**Supplementary Table 1B**). Interestingly, though we were able to match 18,084 T-box sequences with tRNAs, we only identified a tRNA pair for 8.6% of T-boxes with 3′-U specifiers, consistent with observations that 5′-A starting anticodons in bacterial tRNAs are rare (**Supplementary Table 1C**). In these cases, it is possible that these T-boxes are controlled by tRNAs without matching anticodons (e.g. rely on wobble base-pairing) or are using an alternative specifier reading frame when binding anticodons. Alternative reading frames for T-box sequences have previously been observed in at least one experimentally studied system [40].

### T-box specifier usage

Identifying specifier sequences for T-boxes allowed us to interrogate the choice of tRNA anticodon, and therefore the tRNA, that is used for regulation. Analogous to ‘codon usage tables’, which summarize an organism’s codon preference for translating a particular amino acid, **Table 1** depicts a ‘specifier usage table’ generated from 23,497 sequences in the TBDB. For T-boxes of amino acid families that only have two codons (Lys, Asp, Asn, Glu, Gln, His, Tyr, Cys, Phe), a single specifier is preferred in over 85% of corresponding T-box sequences. The choice of specifier is also consistent, with 3′-A and 3′-C always favored over 3′-G and 3′-U respectively. T-boxes of amino acid families decoded by four codons (Gly, Ala, Pro, Val, Thr) have more diversity in specifier usage. Much like the two-codon sets, there is a preference for Val, Thr, and Gly T-boxes to use 3′-A and 3′-C specifiers. Interestingly, Ala and Pro T-boxes display a preference for 3′-U specifiers. T-box families for amino acids with 6 codons (Leu, Ser, Arg) show a 3′-A and 3′-C preference, with an even stronger preference for 3′-C specifiers. In the case of Leu family T-boxes, the CUC specifier is observed in 74% of sequences. For the special case of Ile, 3′-C (AUC) specifier is preferred, though there is potential ambiguity in cases where Ile and Met tRNA share anticodons. The large-collection of sequences has allowed us to reinforce previous observations that the “C-rule” (3′-C in specifiers) is prevalent, while additionally discovering that 3′-A usage is also largely preferred for specific amino-acid classes (**Supplementary Table 1BC**) [23,24,36].

There are possible explanations for the source of specifier-usage bias in T-box riboswitches. First, specifier usage does not follow the same observed patterns of codon usage. In most cases, the preferred specifier is the least preferred codon for the amino acid family. For example in the taxonomical order Bacillales, the Phe UUU codon is used in approximately 70% of cases for translation, but is present in only 1% of Phe T-box specifiers [42]. One possible hypothesis for the specifier-use bias could be attributed to T-boxes favoring interaction with a single tRNA species and disfavoring wobble base-pairing. In the absence of tRNA with 5′-I (Inosine) anticodons, 3′-A and 3′-C anticodons are only decoded by a single tRNA species (5′-U and 5′-C anticodons), whereas 3′-G and 3′-U codons can be decoded by multiple tRNAs (5′-U/C and 5′-A/G anticodons). T-boxes likely co-evolved to be highly specific in their response towards a single tRNA species, which would have been made more difficult if specifier binding is made competitive with two (or more) tRNA species (Watson-Crick basepair vs wobble). Additionally, tRNAs with 5′-A anticodons are not prevalent in bacteria, as the U:G wobble-pair is the preferred mechanism for decoding 3′-U codons [43]. The consensus sequence of the 23,497 T-boxes (**Supplementary Figure 10**) revealed that 5’-NNC-3’ specifiers were preferred overall, being represented at 57.6% of T-box sequences with predicted specifiers.

### Distribution of T-boxes in TBDB by phyla

A majority of the transcriptional T-box collection in TBDB primarily derives from gram-positive phylum Firmicutes (Figure 3) [44]. In particular, sequences in the Bacilli and Clostridia family, within the phylum Firmicutes, constitute the majority of T-box sequences. T-boxes with specifiers matching all the standard 20 tRNA families are observed in Bacilli and Clostridia, and agrees with the prevailing notion that these bacteria use T-boxes as primary regulators for controlling aminoacylation homeostasis. For non-Firmicutes sequences in the TBDB, the diversity and distribution of T-box does not follow the trend within Firmicutes and likely reflect phyla specific usage and adaptation. The other major family represented in the TBDB collection are the Ile translational T-boxes derived from Actinobacteria. Overall the Ile, Trp, Leu, Met and Val families constitute 50% of the T-boxes in TBDB. The diversity of T-boxes in Firmicutes, and relative lack thereof in other phyla, brings to question their origins. The distribution of T-boxes in TBDB conforms to previous observations that T-boxes likely evolved in a common ancestor of Firmicutes and the Actinobacteria, the Chloroflexi, and the Deinococcus-Thermus (DT) group while potentially being distributed across other phyla through horizontal gene transfer (HGT) events [23].

**Fig 3.**
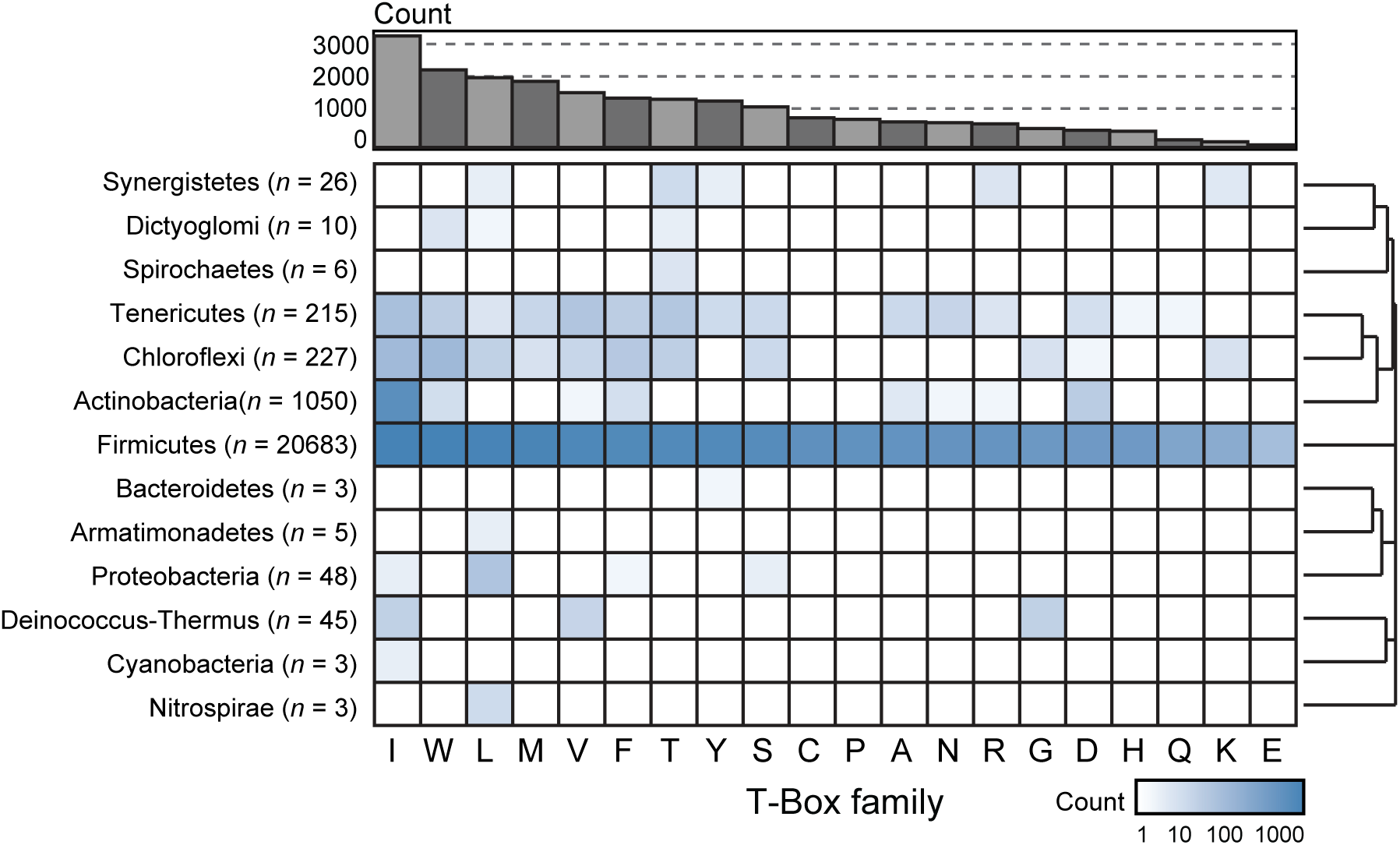
Composition of TBDB by phyla and T-box family. TBDB contains 23,497 T-box sequences from 3,621 different species. The majority of sequences (20,683) in our collection originate from the phyla Firmicutes. T-boxes shown are displayed in decreasing order of abundance in the TBDB collection, with Ile being the most commonly represented and Glu the least. Phyla were clustered by hierarchical clustering using binary distance metric from the Superheat R package [47]. Heatmap displays abundance (total number) of individual T-box families for each phyla in our TBDB collection, using a log_10_ scale to visualize low abundance distribution.

### Conclusion

As riboswitches, T-boxes were the first to be discovered but remain under characterized. Of the known >20,000 sequences, few have been tested for regulatory activity. Currently, T-box research is stymied by the necessity of secondary structure modeling to resolve which tRNA a given T-box binds. Through a comprehensive compilation of sequence information, the TBDB increases access to T-box functional information. The TBDB has aggregated and processed over 23,000 T-box sequences in order to identify structural folds and tRNA binding partners. The TBDB aims to be an approachable hub for the riboswitch community and in future version aims to integrate well with experiments with both natural and engineered T-box riboswitches. As more experiments are being carried out in this area, additional feature annotations will become available - such as predictive models for tandem T-boxes and mutability of synthetic T-boxes.

## Materials and Methods

### Data collection and sequence curation

Transcriptional T-box leader sequences used to generate the database were obtained from a variety of primary sources including RFAM 14.0 database, RibEX database, GeConT3 database, and others [24,35–38]. Redundant sequences were removed from the input dataset before further processing. For T-box leader sequences that did not necessarily contain the terminator (RFAM and GeConT3 sequences) Entrez (NCBI) was used to extend sequences by 50 nt. Extended sequences were then subsequently trimmed to end in penta poly-U (3′-UUUUU). Sequences from these databases that were too short (<100 nt) or too long (>500 nt) were also removed. After structural prediction, sequences with INFERNAL scores under 15 were removed (**Figure S11**). Duplicate T-boxes were removed and a unique ID (TBID) was generated for each T-box.

### Structural prediction of T-box leaders and identification of specifier and T-box sequence

INFERNAL was used to predict the secondary structure of input sequences using the RFAM 14.0 T-box leader sequence covariance model (RF00230.cm) [34]. The secondary structure predicted by the covariance model was then used to predict T-box features. The first stem-loop in the secondary structure was assigned as Stem I. The last bulge within the Stem 1 structure was assigned as the specifier bulge. The specifier sequence was assigned as the bases 2-4 (inclusive) 5′- from the end of the Stem I specifier bulge (**Figure S10C**). The antiterminator was assigned as the last stem-loop predicted by the covariance model. The T-Box sequence was assigned as the first unpaired 5′-UGGN-3′ sequence within the antiterminator bulge, and the antidiscriminator base was assigned as the last base of this sequence.

### Detection of isoleucyl translational T-boxes

A set of 115 unique *ileS* leader sequences was generated by BLAST search using a set of 37 published *ileS* T-box sequences as inputs [8,24]. These sequences were aligned using mlocarna, and the multiple sequence alignment was manually annotated with secondary structure corresponding to the published structure of the *Nocardia farcinica ileS* T-box [17,45]. Next, INFERNAL was used to construct a covariance model while iteratively refining the alignment. The covariance model was then applied to search all NCBI reference genomes in the Actinobacteria phylum (TaxID:1760). The covariance model used to identify translational T-boxes is available in our repository (https://github.com/mpiersonsmela/tbox/blob/master/translational/translational_ILE.cm). Finally, features were annotated using the same method as for transcriptional T-boxes, except that the specifier loop was assigned as the Stem 1 hairpin loop.

### T-box structure refinement

The INFERNAL predicted secondary structure and sequence were processed to remove gaps and truncations. For each entry, truncations were filled from the corresponding FASTA sequence. Gaps in the aligned sequence and their corresponding structural characters in the dot-bracket notation were removed. Pairs between nucleotides and gaps in the INFERNAL predicted structure were unpaired by removing the gap/bracket from the sequence/structure, respectively, and changing the paired bracket to a dot. Energetically unfavorable hairpin loops of 0, 1, or 2 nt were expanded into loops of 3 or greater. Antiterminator structures were further refined by 1) using the INFERNAL antiterminator structure as a soft constraint for RNAfold (ViennaRNA) and 2) using the discriminator and the nucleotides downstream of the antiterminator unpaired using hard constraints [33].

The terminator region was found by searching for the first poly-U sequence (defined as a seven-nucleotide region containing at least five U’s) starting 10 nt after the end of the antiterminator hairpin. Once this sequence was identified, RNALfold was used to generate a list of candidate terminator hairpins between the discriminator region and the poly-U region. Next, this list was filtered to remove hairpins with stem length shorter than 6 nt, as well as hairpins ending more than 2 nt before the poly-U region. The terminator hairpin was chosen as the largest hairpin among this filtered list, and the energy was evaluated using RNAeval. If this search method failed, due to lack of a poly-U region or otherwise, a fallback method simply used RNAfold to find the minimum free energy structure of the sequence from the antiterminator start to the end of the T-box. Energies of terminators predicted by the fallback method were not included in analysis. Terminator energies were not predicted for translational T-boxes.

### Thermodynamic calculations

RNAeval (ViennaRNA) was used to calculate Gibbs free energy (MFE) of predicted secondary structures for terminator and antiterminator sequences [33]. The sequence and associated secondary structures starting at the antiterminator and ending at the poly-U if present were used to calculate ΔG_anti_ (kcal/mol) and ΔG_term_ (kcal/mol). Calculations were performed at 37 °C with default settings, allowing GU pairing. The free energy change between terminator and antiterminator conformations for the T-box ΔΔG_term-anti_ was taken as the difference between ΔG_term_ and ΔG_anti_. Thermodynamic contributions from tRNA binding are not taken into consideration.

### Pairing T-boxes with putative cognate tRNAs

Top specifier sequence predictions were used to identify a putative tRNA family pair for each T-box. To find the sequence of cognate tRNA, Entrez (NCBI) was used to query for all genomic records of the T-box host organism. Genome sequences, either partial or full, were downloaded from RefSeq or GenBank. tRNAscan-SE (Lowe Lab, UCSD) was used to identify all possible tRNAs in each host organism. tRNAs with predicted anticodons matching the specifier were considered matches. For cases where more than one possible tRNA gene was possible, a single tRNA was chosen from among the matching tRNAs for display. tRNA visualization was generated using VARNA [31].

### Identification of downstream gene

The T-box NCBI accession was used to identify the genetic locus of the T-box within its host organism. An Entrez query for features 500 nt downstream of the T-box end was used to retrieve downstream gene/protein description, protein accession, and enzyme commission number if available. The protein accession was then used to obtain additional gene ontology information by querying the appropriate database (Uniprot, KEGG, ENA, or DDBJ).

### Prediction of T-box specifier sequence

The T-box specifier region was assigned as the 1-5 bp (inclusive) 5′-from the Stem I specifier bulge. In these five bases, three possible specifier reading frames (−1, +0, +1) were examined for meeting specifier-match criteria. For each possible specifier, we identify the putative tRNA family (by matching anticodon) that would bind. We then check to see if 1) the predicted tRNA family had a discriminator base that could base pair with the T-box sequence, with wobble allowed, and 2) if the predicted tRNA amino acid family matches the downstream gene ontology. In the case of predicted His T-boxes, discriminators were not used as a criterion for matching, as most mature tRNA^His^ transcripts have a paired discriminator base. For matching T-box sequence with tRNA discriminators, we first searched the host for all tRNAs of given tRNA family and identified which discriminator base that specific host used. In cases where we could not identify matching tRNAs in the host organism, the bacterial discriminator base frequency information was extracted from tRNAviz and utilized in the specifier prediction model [46]. The top specifier was then assigned as the specifier that met most of these conditions, equally weighted, with preference given in the following order: +0 > −1 > +1 specifier reading frames. In cases where more than one specifier was possible, the top specifier was assigned as mentioned but alternative specifier reading frames are also provided in TBDB.

### Benchmarking specifier feature prediction

The feature-annotated T-box leader dataset from Vitreschak *et al.* was used to benchmark the pipeline for structure and feature prediction [24]. Of the initial 698 sequences from the curated input Vitreschak *et al.* dataset, INFERNAL was able to detect T-box leaders in 694 sequences (99.5%). Of these, 621 (89.5%) scored high enough to match both specifier bulge and T-box sequence predictions. Our structure prediction pipeline was able to accurately predict 589 of 621 specifier sequences (94.8%) from the Vitreschak *et al.* dataset [24]. In one additional sequence the appropriate specifier was found in the “−1” specifier frame (0.2%) and in 9 other sequences the correct specifier was found “+1” specifier frame (1.4%) (**Supplementary Figure S10C**). 22 sequences (3.5%) had specifier predictions that did not match the Vitreschak *et al* dataset predictions and include cases where specifier predictions were off by 2 or more nucleotide.

### Visualization of T-box structures

The MFE structure predictions of the refined terminator and antiterminator structures were combined with the INFERNAL prediction structure of the Stem I, Stem III, and other regions, if present [34]. VARNA was used to convert structures from dot-bracket notation to 2D flat image representation [31].

## Supporting information

TBDB_database_field_description

TBDB_database_flatfile

## Data availability

The TBDB is free to access and does not require user registration to use. The database is accessible to browse at https://tbdb.io. All data used to generate TBDB can be accessed for download at https://tbdb.io/download. The full pipeline used to generate entries in TBDB (from FASTA to TBDB entry) is available to download in our repository (https://github.com/mpiersonsmela/tbox/). Additionally, a site repository is also available that contains the front-end of the website.

## Acknowledgements

Funding for this work was provided by U.S Department of Energy grant DE-FG02-02ER63445. Merrick Pierson Smela was supported by an NSF graduate research fellowship. We thank Alex Mijalis, Devon Stork, Erkin Kuru, Anush Chiappino-pepe, and John Aach for their feedback on the database and manuscript.

## Author contributions

Project conceptualization, KN and JAM; Source code, MPS, JAM and THHJ; Database creation JAM and MPS; Data analysis: JAM, MPS, KN and THHJ; manuscript writing and editing, JAM and KN with assistance from MPS and THHJ; Supervision, KN, JAM and GMC.

## Supplementary Results and Discussion

**Table S1.** Sequence composition of the T-box Database **A) Composition of T-box database by tRNA family.**

A) Composition of T-box database by tRNA family.
| tRNA family | # Sequences | # Sequences<br>(w/ tRNA) |
| --- | --- | --- |
| ALA | 759 | 159 |
| ARG | 706 | 321 |
| ASN | 753 | 635 |
| ASP | 510 | 451 |
| CYS | 878 | 836 |
| GLN | 226 | 217 |
| GLU | 51 | 43 |
| GLY | 556 | 523 |
| HIS | 476 | 425 |
| ILE | 3320 | 2769 |
| LEU | 2099 | 1481 |
| LYS | 171 | 158 |
| MET | 1951 | 1783 |
| SUP | 9 | 0 |
| PHE | 1483 | 1321 |
| PRO | 837 | 314 |
| SER | 1225 | 895 |
| THR | 1430 | 1023 |
| TRP | 2337 | 2133 |
| TYR | 1393 | 1266 |
| VAL | 1642 | 1331 |
| NONE | 685 | 0 |

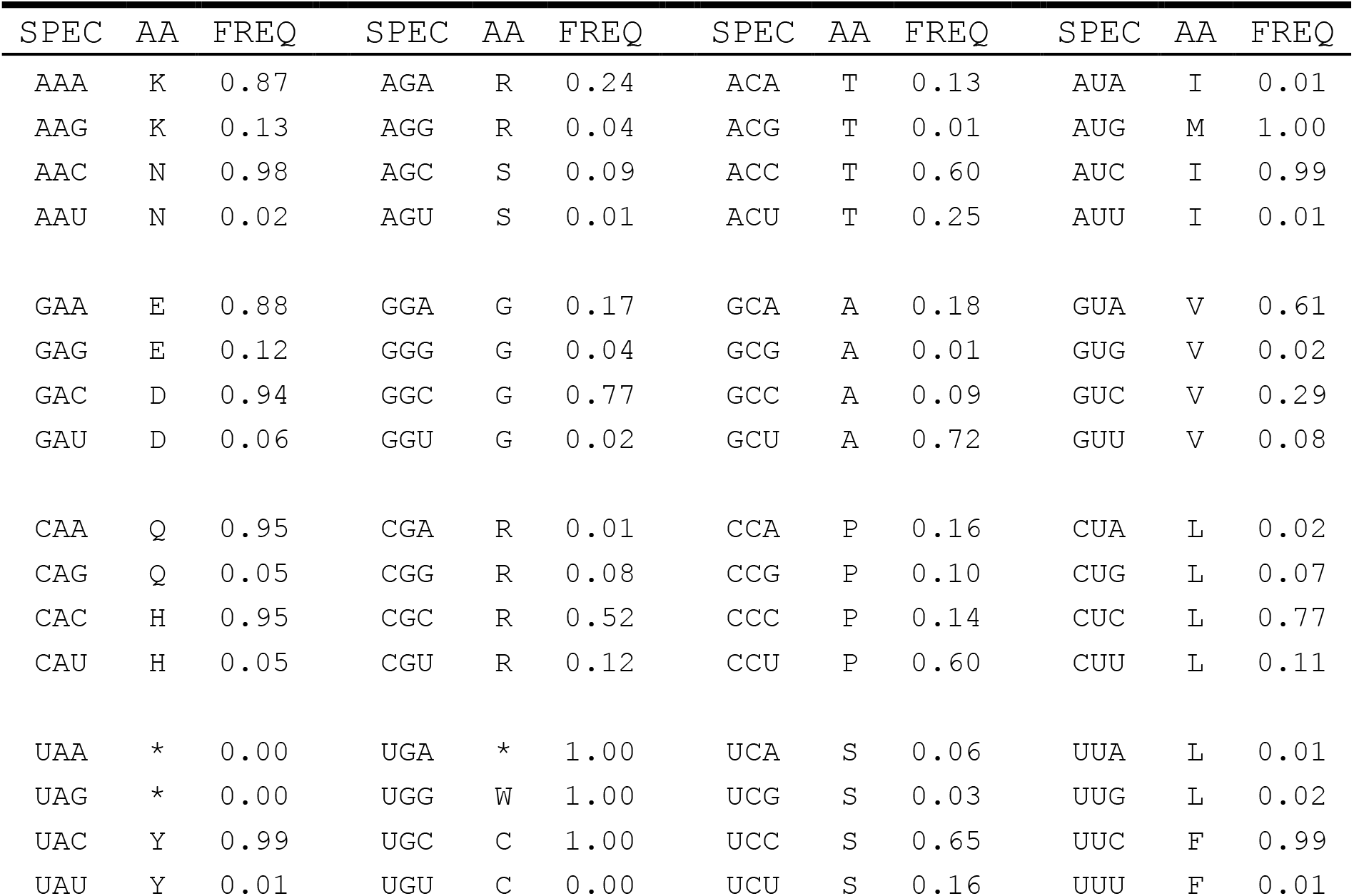
B) Composition of T-box database by predicted specifier sequence. ‘SPEC’ column indicates predicted specifier sequence (5′-3′), ‘SEQ’ shows total number of sequences with predicted specifier in the TBDB, and ‘w/ tRNA’ shows the number of sequences where tRNAscan-SE found a matching tRNA in the host genome.

C) Composition of T-box database by 3 ′-base of predicted specifier sequence.

| Specifier | # Sequences | # Sequences<br>(w/ tRNA) | % |
| --- | --- | --- | --- |
| NNA | 2290 | 2006 | 87.6% |
| NNU | 2191 | 189 | 8.6% |
| NNC | 13538 | 11602 | 85.7% |
| NNG | 4793 | 4287 | 89.4% |

**Table S2.** Summary of RNA folding energies for sequences in the database. Mean and standard deviation Δ*G_anti_*, Δ*G_term_*, ΔΔ*G_term-anti_* for all sequences in the database, grouped by tRNA family

| tRNA family | $\Delta G_{anti}$ (kcal/mol) | | $\Delta G_{term}$ (kcal/mol) | | $\Delta\Delta G_{term-anti}$ (kcal/mol) | |
| --- | --- | --- | --- | --- | --- | --- |
| | $\mu$ | $\sigma$ | $\mu$ | $\sigma$ | $\mu$ | $\sigma$ |
| ALA | -7.1 | 2.2 | -16.2 | 7.5 | -9.1 | 6.7 |
| ARG | -5.9 | 2.2 | -17.4 | 5.4 | -11.5 | 4.4 |
| ASN | -7.2 | 2.2 | -20.0 | 6.3 | -12.9 | 5.6 |
| ASP | -7.5 | 2.6 | -19.8 | 7.7 | -12.6 | 6.6 |
| CYS | -5.7 | 1.8 | -19.1 | 5.5 | -13.5 | 5.3 |
| GLN | -8.1 | 2.8 | -21.8 | 5.8 | -13.6 | 5.0 |
| GLU | -6.9 | 1.8 | -19.5 | 5.7 | -12.8 | 4.8 |
| GLY | -9.1 | 2.9 | -24.1 | 12.0 | -15.0 | 10.5 |
| HIS | -6.1 | 2.4 | -20.0 | 5.2 | -13.9 | 4.6 |
| ILE | -11.1 | 7.9 | -19.2 | 10.2 | -12.2 | 8.2 |
| LEU | -8.0 | 3.1 | -20.5 | 7.3 | -12.5 | 6.2 |
| LYS | -7.4 | 2.2 | -21.0 | 5.9 | -13.6 | 4.8 |
| MET | -5.8 | 2.8 | -20.4 | 6.9 | -14.7 | 6.8 |
| PHE | -6.7 | 2.3 | -17.4 | 5.7 | -10.7 | 5.5 |
| PRO | -7.1 | 2.0 | -19.1 | 6.3 | -12.1 | 5.9 |
| SER | -7.4 | 2.2 | -19.2 | 6.8 | -11.8 | 6.4 |
| THR | -6.8 | 2.5 | -19.4 | 5.7 | -12.6 | 5.1 |
| TRP | -6.7 | 2.5 | -19.9 | 6.4 | -13.4 | 6.1 |
| TYR | -6.6 | 2.0 | -18.5 | 5.2 | -12.0 | 5.0 |
| VAL | -5.5 | 2.3 | -18.6 | 8.2 | -13.1 | 6.9 |

**Fig S1.**
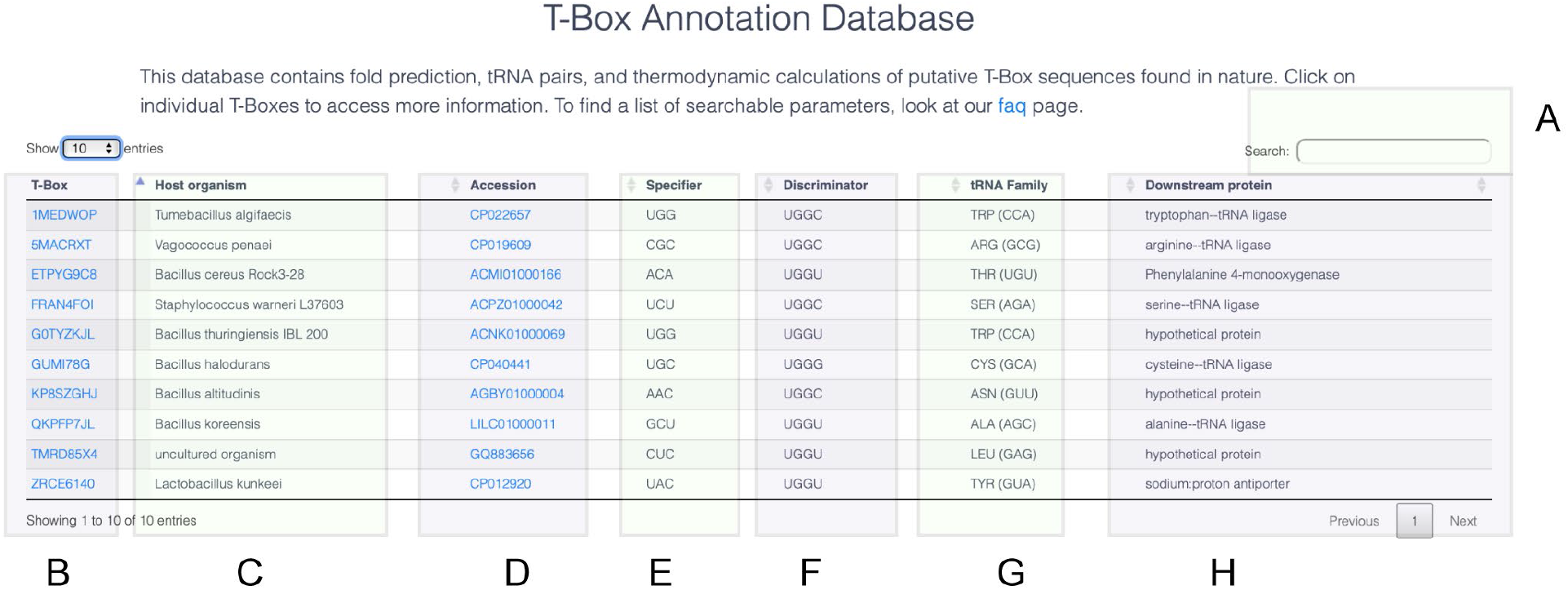
The TBDB interface – Database table. A) Entries into the TBDB are searchable by a variety of fields including sequence, specifier, tRNA family, amino acid family, discriminator bases, and downstream protein. B) The T-box column displays name of T-box by TBDB unique ID. Clicking on an individual ID brings users to a T-box entry page, where additional information is displayed. C) The ‘Host Organism’ column displays species from which the T-box input sequence was obtained from. D) Clicking on genomic accession in the ‘Accession’ column brings users to an NCBI genetic locus entry page for a T-box. E) The ‘Specifier’ column shows predicted specifier sequence (5′-3′) for given T-box. F) The ‘discriminator’ column shows the T-box sequence, or anti-acceptor arm. G) ‘tRNA Family’ indicates the type of tRNA, alongside the corresponding amino acid, with anticodon matching the specifier sequence. H) A description of the protein found downstream of the T-box is provided in the ‘Downstream protein’ column.

**Fig S2.**
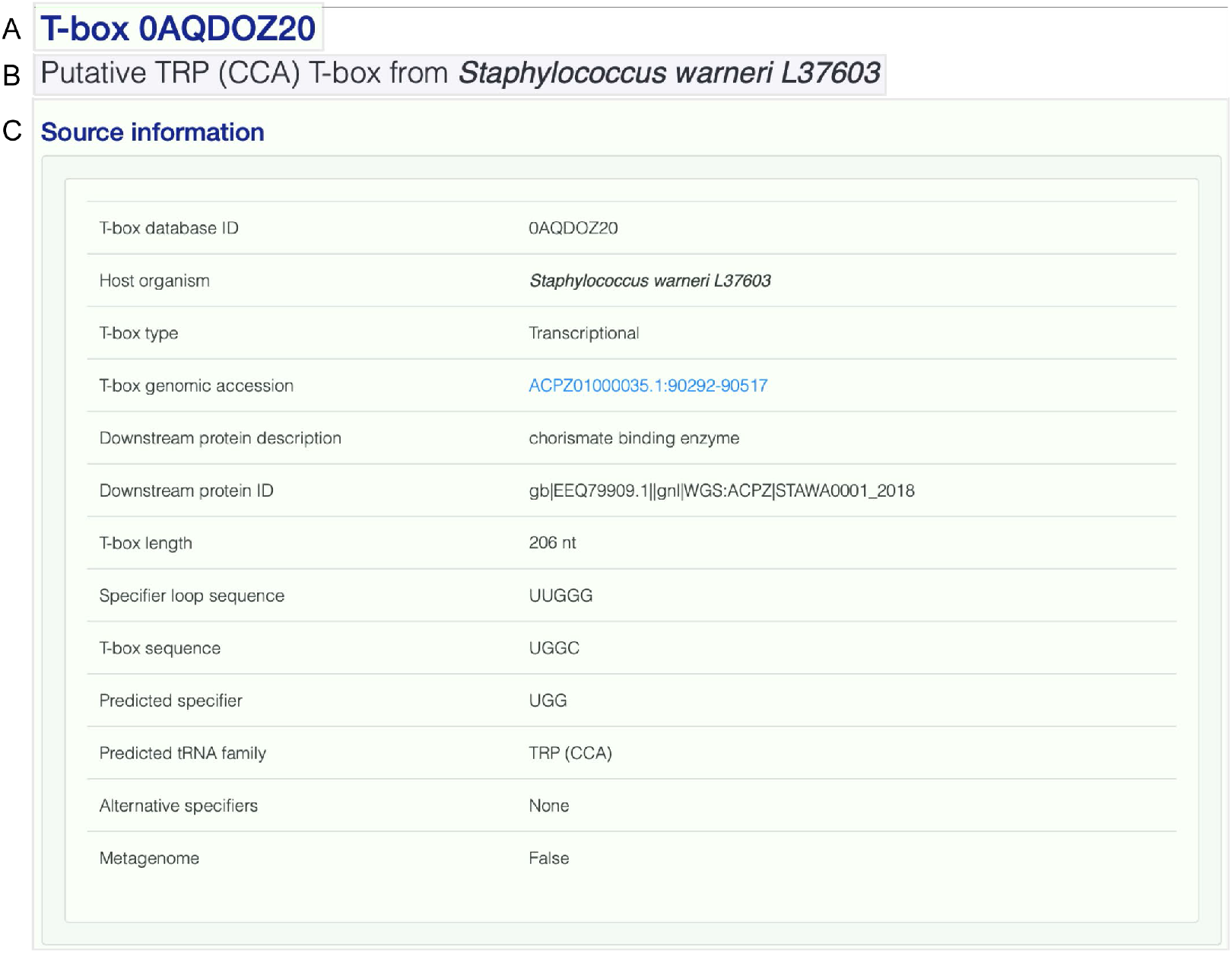
The TBDB interface – Entry title and source information. A) Entries into the TBDB are titled using the TBDB unique ID. B) A brief description of the T-box is also provided, and includes tRNA family displayed as amino acid (anti-codon), and host organism where T-box input sequence was found. C) The ‘Source Information’ panel provides a summary of important T-box features.

**Fig S3.**
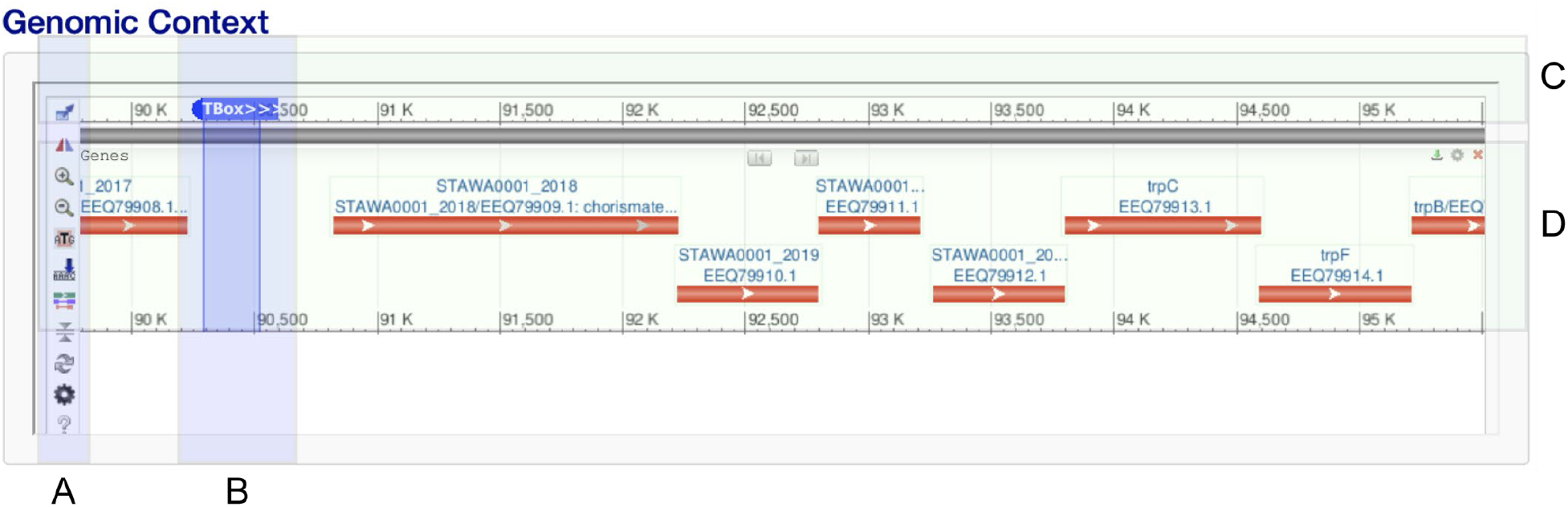
The TBDB interface – Genomic context viewer. An integrated NCBI graphical display of T-box genomic region can be used to interrogate genes or operons controlled by T-box riboswitches from host organism. Here, a Trp T-box is shown upstream of a tryptophan biosynthesis operon. A) Browsing menu allows users to change a variety of settings for the NCBI graphical viewer. B) T-box input sequence is highlighted in blue, with arrows indicating the direction, (+) or (−) strand of T-box transcription. C) Genomic region is shown as the top gray bar, with numbers indicating genomic locus. D) Genes are provided in the center panel and are annotated as provided by NCBI.

**Fig S4.**
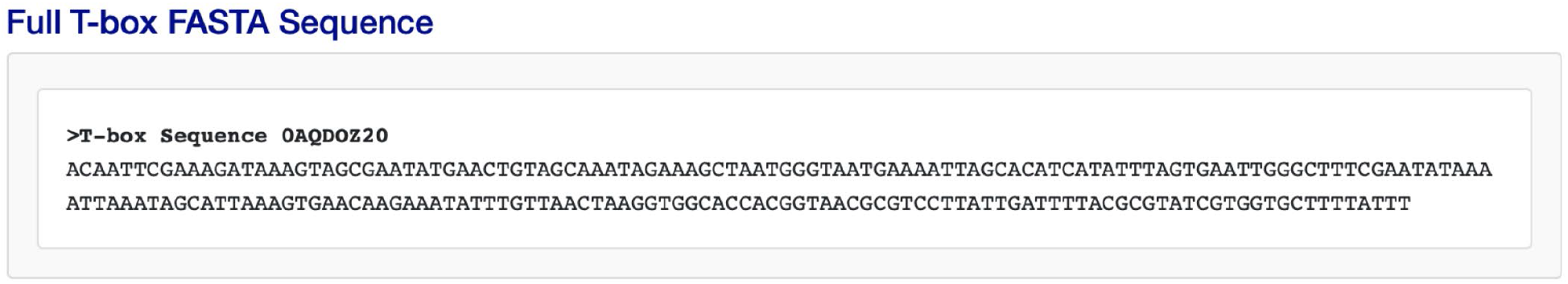
The TBDB interface – T-box FASTA sequence. The FASTA sequence of T-box, starting from Stem I and ending at the terminator, is provided as a panel. Sequence header uses the TBDB unique ID.

**Fig S5.**
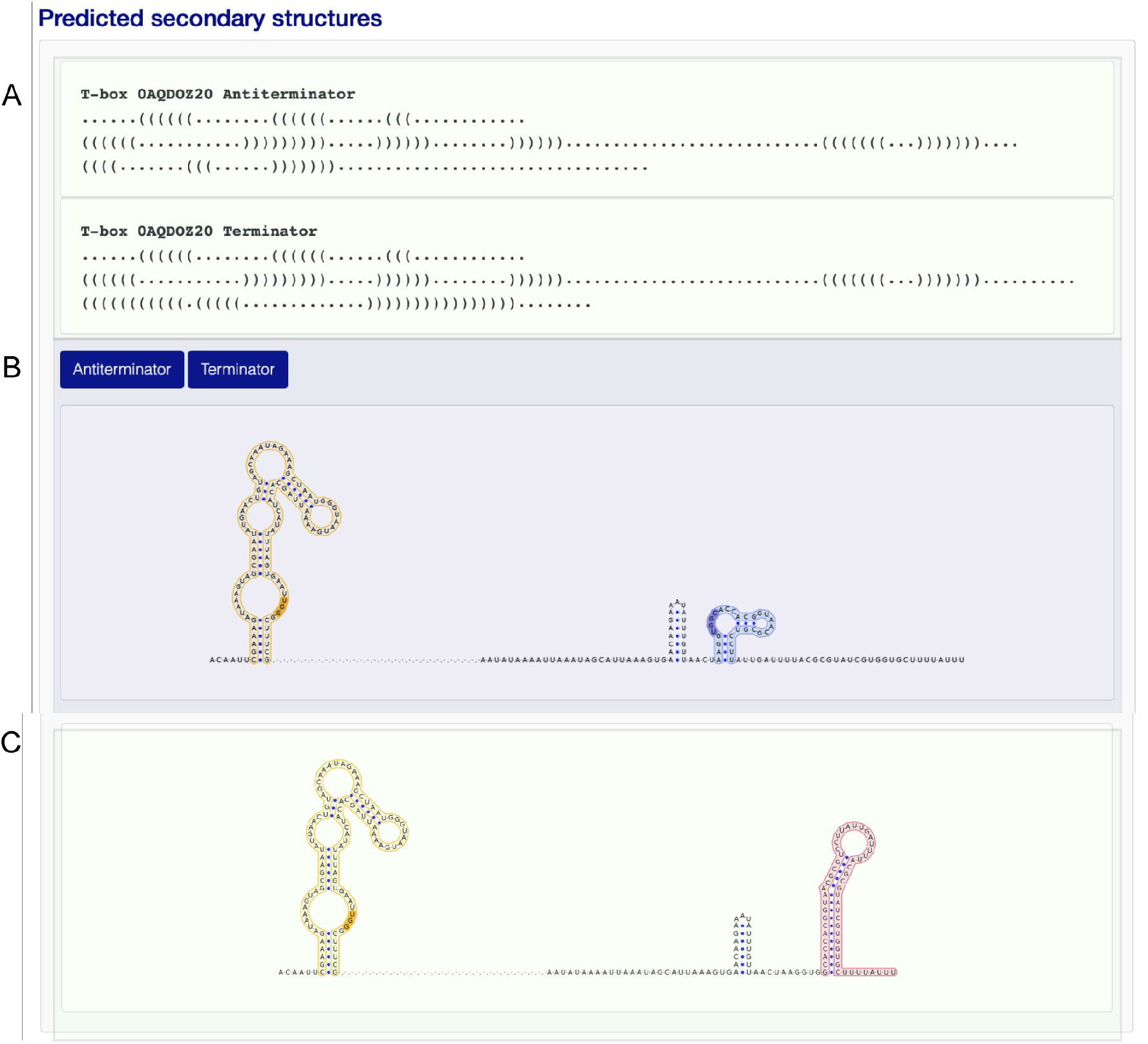
The TBDB interface – Predicted secondary structure visualization. Secondary structures for T-boxes, as predicted by folding and refinement, are provided as dot-bracket structures for antiterminator and terminator. The dot-bracket representations can be used alongside the FASTA sequence provided in the “Full T-box FASTA sequence” panel to generate custom 2-D representations of T-boxes. Headers use TBDB unique ID for each T-box. A) Two secondary structure representations are provided, if available, for each T-box sequence. For transcriptional T-boxes, the ‘Antiterminator’ option shows the antiterminator fold while the ‘Terminator’ option shows the terminator fold representation. Sequences are displayed using VARNA flat representation for B) antiterminator and C) terminator conformations. Gaps are inserted after the Stem I structure to improve clarity of display and prevent overlaps between the stem-loops in visualization. Important features are highlighted as follows: Stem I (light yellow), specifier bases (orange), antiterminator (light blue), anti-acceptor arm (blue), and terminator (red). Note that terminator representations are not displayed for sequences where local fold using ViennaRNA failed to find an MFE terminator structure.

**Fig S6.**
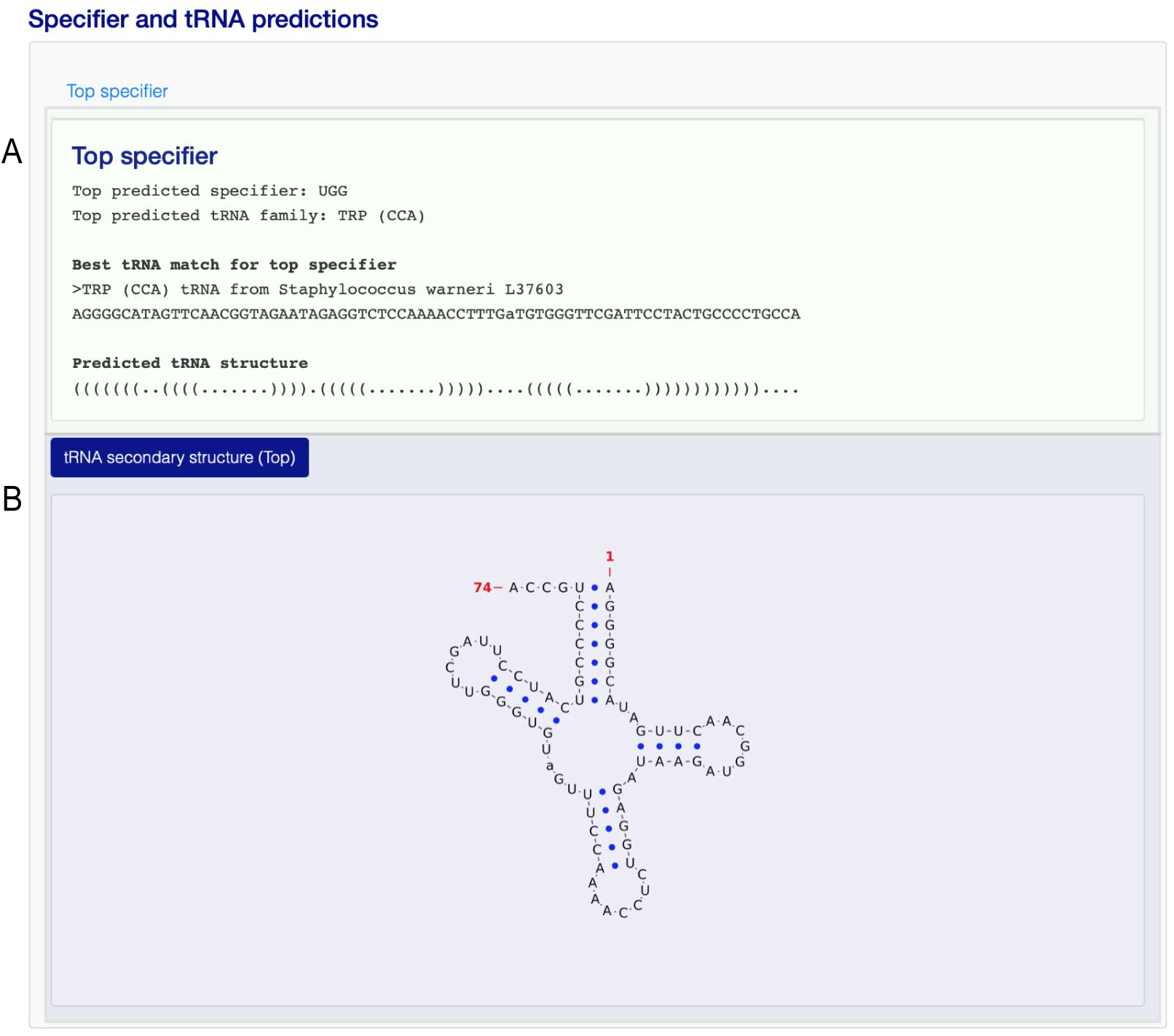
The TBDB interface – tRNA scan results. Results from tRNA scan for a matching tRNA from host organism are provided in the ‘tRNA prediction’ panel. tRNAscan-SE was used on the host organism to identify tRNA with anticodons matching the specifier sequence of the given T-box. A) The sequence and structure, in dot-bracket notation, for the top tRNA match is shown. FASTA header displays tRNA family and host organism. If no tRNA was found in the host, a null result will be displayed. If there was ambiguity in deciding the reading frame, results for alternative reading frames are also displayed. B) For T-boxes where a matching tRNA from native host was found, secondary structure is displayed as a 2-D VARNA generated image.

**Fig S7.**
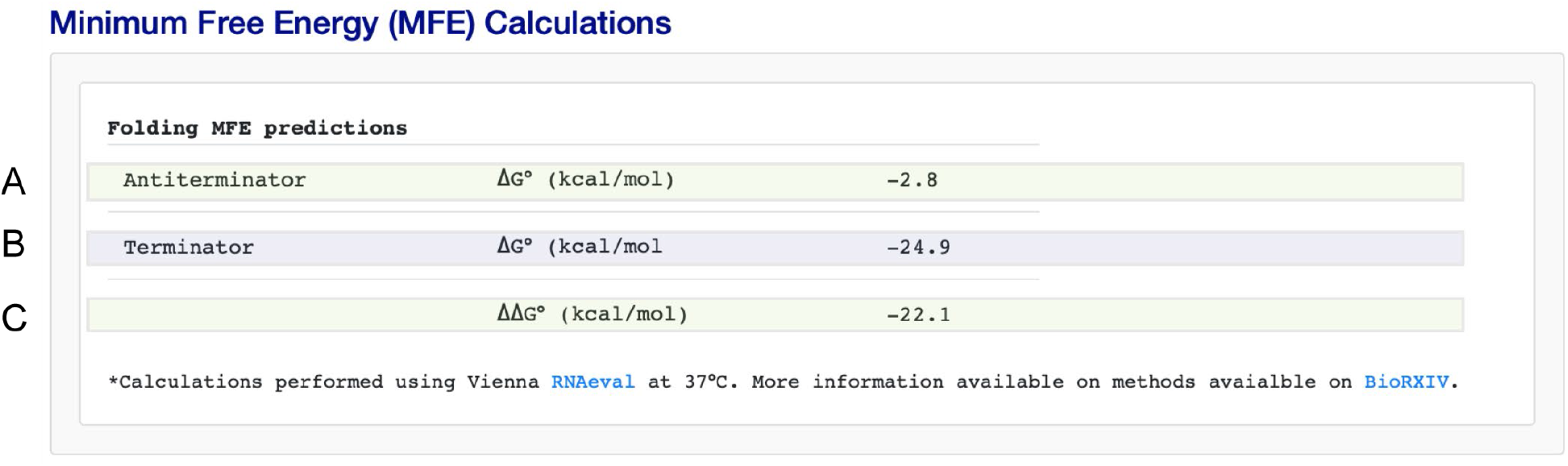
The TBDB interface – MFE predictions. Gibbs free energies from MFE predictions (Vienna RNA) are provided for A) antiterminator structure, B) terminator structure. C) Displays the difference in MFE between terminator and antiterminator conformations.

**Fig S8.**
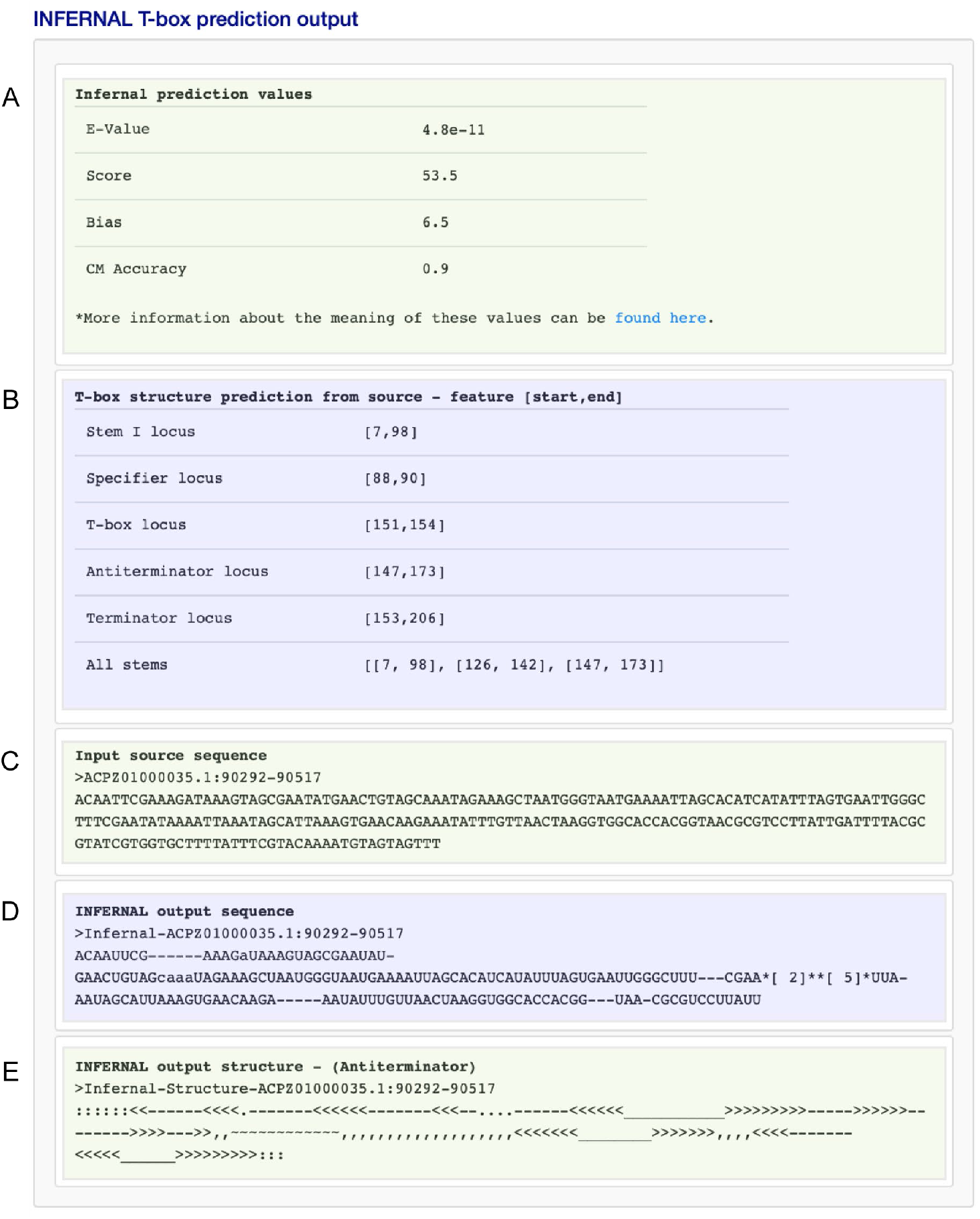
The TBDB interface – INFERNAL prediction output. Information used to predict T-box structures using INFERNAL is provided in the ‘Infernal T-box prediction output’ panel [34]. A) INFERNAL quality score metrics (E-Value, score, bias, and covariance model accuracy (CM)) are provided. B) Feature identification output, using input sequence of T-box, gives the position information for Stem I, antiterminator, terminator, and specifier. All stems found using the INFERNAL model are provided as ‘All Stems’. Loci are shown in [index start, index end] format. C) Input sequence (as obtained from source database) that was used for T-box predictions is shown in FASTA format. Header indicates host organism genomic accession with locus start-end. D) INFERNAL output sequence is provided in FASTA format, as well as E) INFERNAL output structure.

**Fig S9.**
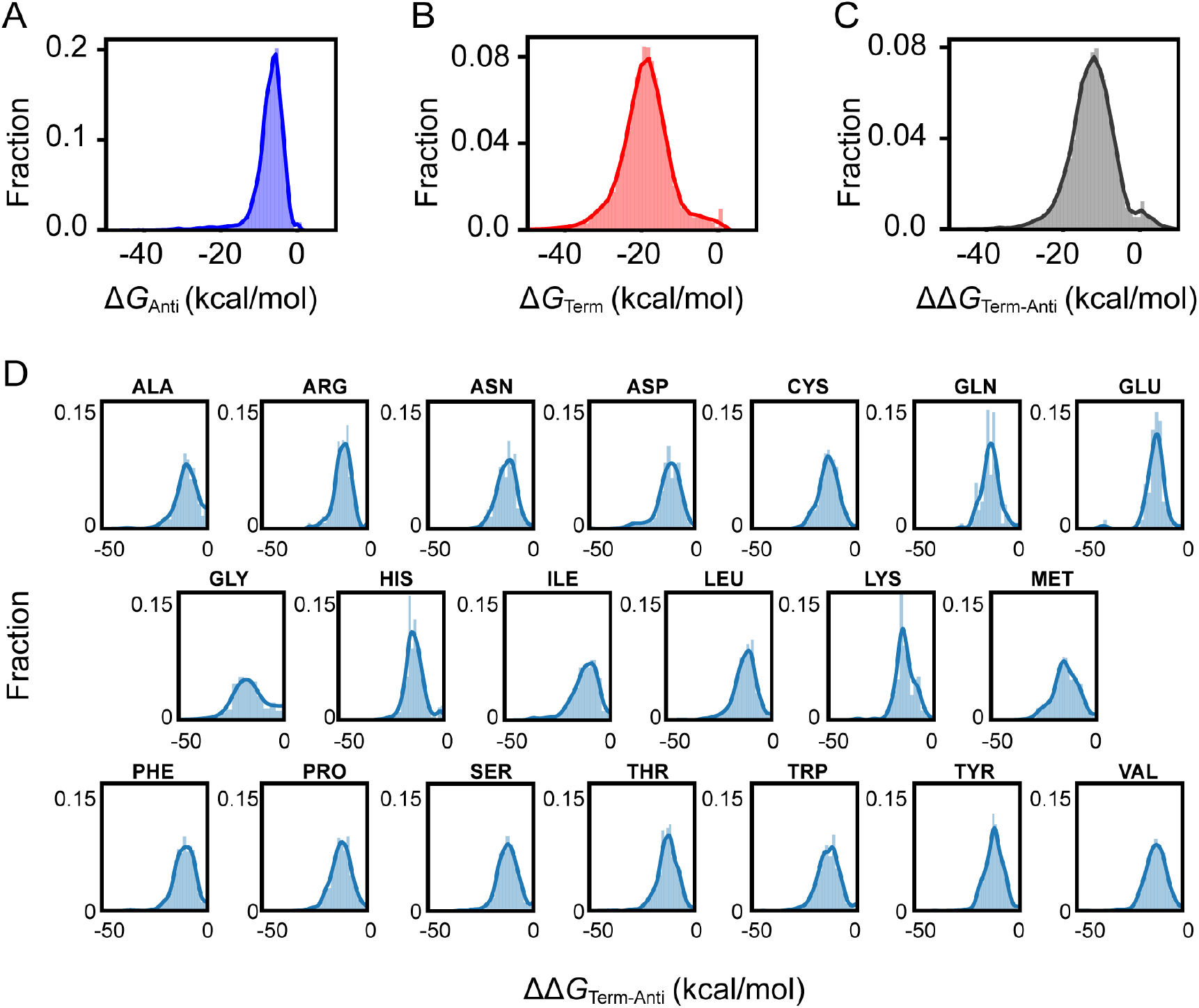
Distribution of minimum free energy structures in the TBDB. A) Distribution of sequences by Gibbs free energy of folding for antiterminator structure (Δ*G_anti_*, kcal/mol). B) Distribution of sequences by Gibbs free energy of folding for terminator structure (Δ*G_term_*, kcal/mol). C) Distribution of sequences by change in Gibbs free energy between terminator and antiterminator conformation (ΔΔ*G_term-anti_*, kcal/mol). D) Density plots showing distributions for change in Gibbs Free Energy (ΔΔG_*term-anti*_) for T-boxes, sorted by tRNA family.

**Fig S10.**
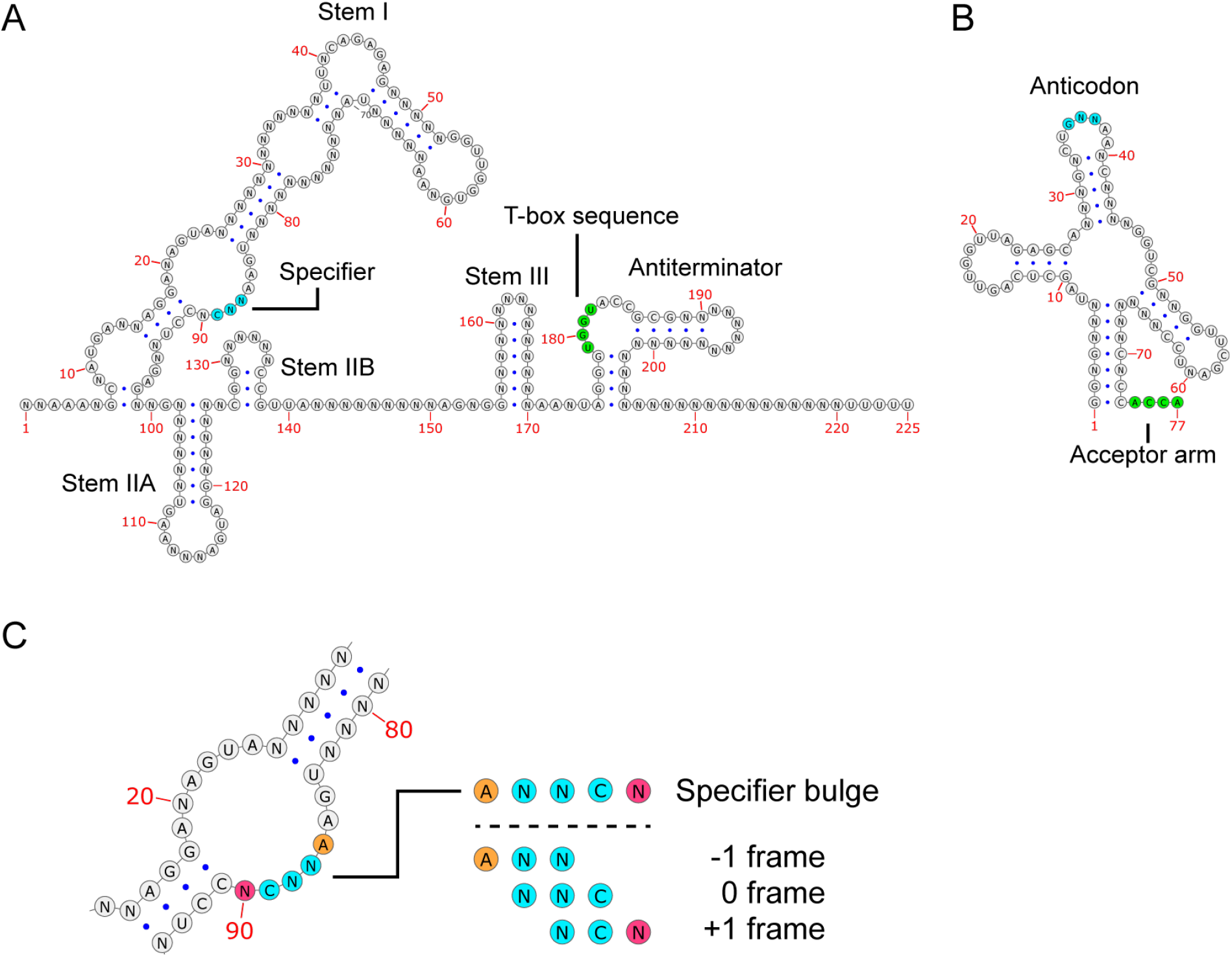
Consensus sequence and structure for T-box and tRNA families in the TBDB. A) T-box consensus sequence from the TBDB with important structural features highlighted. Specifier sequence is highlighted in blue, while the T-box sequence is highlighted in green. B) Consensus tRNA sequence for tRNAs that match with T-boxes in the TBDB. Anticodon is highlighted in blue, while the acceptor arm is highlighted in green. Sequences and structures were generated using cmalign and trimmed using trimal [48]. ‘N’ bases are shown where no base was present in >50% of sequences. C) Three specifier reading frames within the specifier bulge are considered in model decision.

**Fig S11.**
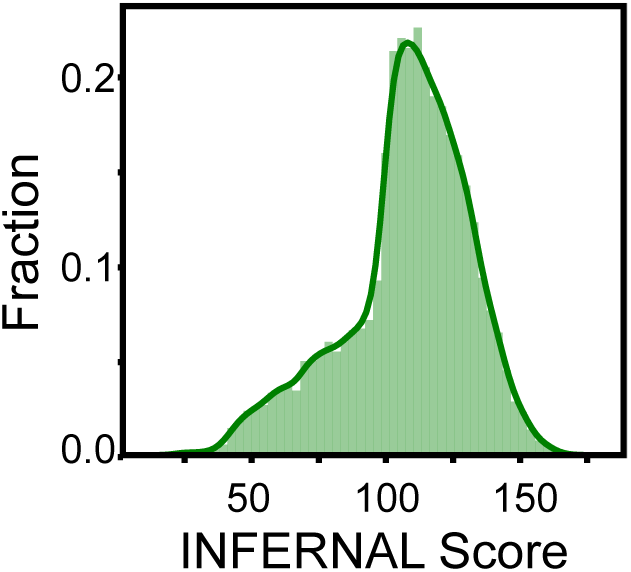
INFERNAL score density plot for sequences in the TBDB. The INFERNAL score is a measure of how well the sequence matches the covariance model and corresponds to the log-odds ratio of the hit sequence relative to random sequence. Sequences scoring <15 (equivalent to an E-value of >0.05) were removed from the database.

## Notes

https://tbdb.io

https://github.com/mpiersonsmela/tbox/

### Competing Interest Statement

This project has been funded by DOE grant DE-FG02-02ER63445. Dr. Church is a founder of the following companies in which he has related financial interests: ReadCoor; EnEvolv; and 64-x. For a complete list of financial interests of Prof. George Church, see also http://arep [dot]med[dot]harvard[dot]edu/gmc/tech[dot]html

